# Heterocyclic sterol probes for live monitoring of sterol trafficking and lysosomal storage disorders

**DOI:** 10.1101/267948

**Authors:** Jarmila Králová, Michal Jurášek, Lucie Krčová, Bohumil Dolenský, Ivan Novotný, Michal Dušek, Zdeňka Rottnerová, Michal Kahle, Pavel Drašar, Petr Bartůněk, Vladimír Král

**Author notes:** Corresponding Author: Jarmila Králová, CZ-OPENSCREEN, Institute of Molecular Genetics of the ASCR, v.v.i. Vídeňská 1083, 142 20 Prague 4, Czech Republic.

## Abstract

The monitoring of intracellular cholesterol homeostasis and trafficking is of great importance because their imbalance leads to many pathologies. Reliable tools for cholesterol detection are in demand. This study presents the design and synthesis of fluorescent probes for cholesterol recognition and demonstrates their selectivity by a variety of methods. The construction of dedicated library of 14 probes was based on heterocyclic (pyridine)-sterol derivatives with various attached fluorophores. The most promising probe, a P1-BODIPY conjugate FP-5, was analyzed in detail and showed an intensive labeling of cellular membranes followed by intracellular redistribution into various cholesterol rich organelles and vesicles. FP-5 displayed a stronger signal, with faster kinetics, than the commercial TF-Chol probe. In addition, cells with pharmacologically disrupted cholesterol transport, or with a genetic mutation of cholesterol transporting protein NPC1, exhibited strong and fast FP-5 labeling in the endo/lysosomal compartment, co-localizing with filipin staining of cholesterol. Hence, FP-5 has high potential as a new probe for monitoring cholesterol trafficking and its disorders.

**Significance statement:** Cholesterol is a vital steroid molecule with many important functions in animal cells. Although its dysregulation is associated with an expanding list of clinically important pathologies, the study of its role is limited by a lack of reliable tools for live intracellular monitoring. This study demonstrates the applicability of a novel class of heterocyclic sterol probes. These probes exhibit fast cellular uptake with effective fluorescence labeling of sterol species in a variety of living cells, without a need for artificial carriers. When applied to Niemann-Pick disease type C1 cells, they identified massive accumulation of cholesterol in the endosome/lysosome compartment. Thus, several probes from the same series can also be used for visualizing lysosomal storage disorders and sterol transporting pathologies.

## Introduction

Cholesterol is a fundamental component of the plasma membrane in animal cells. It controls membrane structural integrity and fluidity, and modulates the activity of various membrane proteins. Membrane physical properties are shaped by interactions of the cholesterol hydroxy group with the polar head groups of membrane phospholipids and sphingolipids, while the bulky steroid and hydrocarbon chain are embedded within the membrane, alongside the nonpolar fatty-acid chains of other lipids.

Cholesterol affects many physiological functions. It reduces the permeability of the plasma membrane to neutral solutes, hydrogen ions, and sodium ions. Within the cell membrane, cholesterol is involved in the formation of invaginated caveolae and clathrin-coated pits, including caveolae-dependent and clathrin-dependent endocytosis. Moreover, cholesterol plays an essential role in the regulation of multiple signaling pathways. It also induces membrane packing in specific microdomains (lipid rafts) of the plasma membrane and provides a platform for a variety of membrane-associated signaling proteins (1). Importantly, cholesterol is also a starting material for the synthesis of steroid hormones (2), bile acids and vitamin D.

Cholesterol is synthesized *de novo* from acetyl-CoA in the endoplasmic reticulum or comes from dietary sources (3, 4). Exogenous cholesterol is acquired through the uptake of low density lipoprotein (LDL) molecules by LDL-receptor mediated endocytosis (5). Biosynthesis, as well as uptake of cholesterol from plasma *via* circulating lipoproteins, is strictly regulated. This regulation involves several feedback loops that ensure the exact amount of cholesterol the cells need for their physiological function. Cholesterol homeostasis in cells is maintained by several mechanisms, including cellular uptake, synthesis, storage and efflux (3, 4). In addition, an involvement of sterol sensing polytopic membrane protein, Scap, that functions as a molecular machine to control the cholesterol content of membranes in mammalian cells has been demonstrated (6). Regarding the distribution, cholesterol moves rapidly between intracellular organelles *via* vesicular trafficking and nonvesicular pathways (3, 7). Excess cellular cholesterol is converted to cholesteryl esters by the enzyme acyl-coenzyme A:cholesterol acyltransferase (ACAT) or is removed from a cell by cellular cholesterol efflux at the plasma membrane (8).

Imbalance in cholesterol homeostasis leads to pathological processes of atherosclerosis, and deregulated cholesterol trafficking is involved in the pathogenesis of neurodegenerative diseases including Niemann-Pick’s disease type C (NPC), Alzheimer’s disease (AD), Parkinson’s diseases (PD), and possibly Huntington’s disease (HD) (9-11). There is also evidence for involvement in steatohepatitis (12, 13).

Due to the critical importance of cholesterol in so many processes, it is fundamental to obtain insight into cholesterol trafficking pathways and kinetics, which have not been fully elucidated yet (14). Filipin, a fluorescent cholesterol-binding polyene antibiotic, is often used for visualization of cellular cholesterol. However, the paraformaldehyde fixation required for filipin labeling itself compromises the morphology of the plasma membrane, and may reorganize membrane components. It is also possible that some intracellular membranes are not as accessible to filipin as the plasma membrane (15) and furthermore, the specificity of filipin for cholesterol is questionable (16).

The development of suitable cholesterol probes for analysis in living cells is a great technical challenge and the search for faithful tools is still ongoing. The candidate molecules have to fulfil requirements for very close biophysical and biochemical resemblance with cholesterol, but at the same time, to display good fluorescence properties. Over recent years, a variety of fluorescent and photoreactive cholesterol probes have been developed (17). So far, mainly two types of fluorescent cholesterol analogues have been used. First, the intrinsically fluorescent mimics of cholesterol, such as cholestatrienol (CTL) and dehydroergosterol (DHE), which exhibit minimal chemical alteration compared to cholesterol, provide low fluorescence signals in the UV region of the spectrum. Second, synthetic cholesterol analogues labeled with fluorophores such as NBD-cholesterol, Dansyl-cholesterol, BODIPY-cholesterol, and fluorescent PEG-cholesterol are often used for microscopic imaging of sterols (17-20). Recently, the click chemistry method based on alkyne cholesterol and oxysterol analogs has emerged as a promising strategy combining benefits of both strong fluorescence and minimal alteration of molecular structure (21, 22). Alternatively, fluorescently labeled cholesterol-recognizing peptides can be used for imaging of lipid domains on plasma membranes (23, 24).

Our approach to this topic was the preparation of new probes comprising heterocyclic steroids conjugated with various fluorophores. Herein, we show the impact of the fluorophore itself and its position on the specificity of binding, localization and trafficking inside the cells. The precursor P-1 with BODIPY linked to the pyridyl moiety (FP-5) exhibited high cholesterol- and cholesterol acetate-binding specificity in spectroscopic studies and fast labeling of cholesterol rich compartments in cellular studies.

## Results

### Molecular design and synthesis of fluorescent probes (FP) with selectivity for sterols

A small dedicated library of fluorescent probes, recognizing cholesterol and cholesterol esters, was synthesized using a multi-block approach. Initially, the steroid-binding unit (steroid skeleton) was substituted with a heterocycle (pyridine) to generate precursors P1 and P2 (Fig. 1), which were then attached to various fluorophores (*SI Appendix*, Figs. S1-S3), giving rise to probes FP-1 – FP-14 (Fig. 1). Importantly, the used precursor P1, is known as abiraterone acetate (brand name Zytiga), which inhibits androgen biosynthesis and is used for treating prostate cancer (25), with documented pharmacokinetics (26). The fluorophores attached on a precursor P1 *via* 3-hydroxy group produced fluorescent probes FP-1 – FP-4 or on pyridyl group *via* quaternization generated probes FP-5–8 (Fig. 1, *SI Appendix*, Fig. S3A). Alternatively, in probes FP-9 and FP-10, the fluorescent group attachment was realized by substitution of P1 on both sites: 3-OH and pyridyl (Fig. 1, *SI Appendix*, Fig. S3A). Another set of fluorescent probes was generated from precursor P-2, where a variety of fluorophores were attached *via* quaternization of the pyridyl group: FP-11–14 (Fig. 1, *SI Appendix*, Fig. S3B). Detailed synthetic protocols and full characterization of all probes are available in *SI Appendix,* 1.Chemistry.

The potential of heterocyclic probes for sensing sterol structures and possible mechanisms of molecular interactions was suggested by X-ray analysis of unique cocrystal of a model sterol (lithocholic acid) with the P1 precursor (*SI Appendix*, 2. Crystallography, Fig. S4, S5). The analysis revealed that the major driving forces for intermolecular bonding of a precursor skeleton and the targeted steroid in an aqueous environment were van der Waals interactions. The hydrophobic effect in aqueous media was a major contributor to cocrystal formation, but possible hydrogen bonding interactions cannot be excluded. The spatial arrangement of the two steroids in cocrystal structure is in an alpha-beta face orientation (*SI Appendix*, Fig. S4, S5; Tables S2-S3)

### Cellular uptake of fluorescent probes

Fluorescent probes for application in cellular studies must be soluble under physiological conditions, and their molecular structure must favour cellular uptake. Therefore, the screen of novel probes was focused on cellular uptake first. Based on fluorescence recorded in U-2 OS cells at various times during incubation (0.5, 8 and 24 h), probes could be divided into several groups as shown in Table 1, *SI Appendix*, Fig. S7: i) exhibiting none or very poor cellular uptake FP-3,4, FP-9, FP-12–14, ii) exhibiting low or mild uptake after longer incubation (8-24 h) FP-1, FP-6, FP-8, iii) probes precipitating in medium with gradual penetration into cell membranes FP-2, FP-10, FP-11, BODIPY-Cholesterol available under commercial name TopFluor-Cholesterol (TF-Chol), iv) exhibiting strong and fast uptake FP-5, FP-7. From these results it can be inferred that the position and structure of fluorophore have a strong impact on cellular uptake. Probes FP-5 – FP-8 with BOPIPY fluorophore connected to pyridyl group *via* quaternization displayed high cellular uptake, while probes FP-1– FP-4 with connection *via* 3-hydroxy group were taken less readily. Furthermore, probes with extended conjugated BODIPY fluorophore (red BOPIPY) FP-3, FP-6, FP8, pyrene FP-4 or coumarin fluorophores FP-1 were internalized less effectively than probes with green BODIPY (FP-2, FP-5, FP-7). In addition, probes FP-9 and FP-10 with the fluorescent group attached on 3-OH and pyridyl sites of P1 and probes FP-11–14 with fluorophores attached *via* quaternization of pyridyl group on precursor P-2 formed visible aggregates (*SI Appendix*, Fig. S7- shown by arrowheads) and enter cells only slowly or not at all. Commercial TF-Chol probe also had a tendency to form aggregates and penetrated cells slowly.

### Unique features of the FP-5 probe

The screening of 14 novel fluorescent probes revealed remarkable properties of FP-5; therefore, this probe is described in detail. FP-5 exhibited fast intracellular uptake, and strong staining of cholesterol rich membranes. In addition, UV-VIS analysis revealed a strong interaction of FP-5 with both cholesterol and cholesterol acetate in aqueous medium. This interaction was accompanied by obvious spectral changes (*SI Appendix*, 3. Spectral analysis, Fig. S6A, B). The absorbance changes were observed at several wavelengths as a function of cholesterol/FP-5 concentration ratio (*SI Appendix*, Fig. S6C, D). Association constants (*log K*) for a 1:1 complex of FP-5:cholesterol and FP-5:cholesterol acetate averaged 5.99 and 6.6, respectively. For 2:1 complexes the values were 11.3 and 12.8, respectively. These values indicated strong binding between partners (*SI Appendix,* Table S4).

Fluorescence spectroscopy analysis revealed a very high fluorescence intensity of FP-5 itself, which rapidly decreased in the presence of sterols (cholesterol, cholesterol acetate) in the aqueous medium due to the formation of complexes (*SI Appendix*, Fig. S6 E, F; Table S5). This is consistent with UV-Vis spectral analysis, suggesting effective binding of selected sterol derivatives with probe FP-5.

Another requirement of probe suitability for live monitoring is its compatibility with cell viability and growth. To establish how FP-5 influences cell growth, we monitored cell proliferation in the presence of increasing concentrations of FP-5 in various cell lines by the IncuCyte live-cell imaging system. Proliferation was monitored by analysing cell counts over one week. U-2 OS cells did not show growth inhibition under 10 μM concentration (*SI Appendix*, 4. Cellular studies, Fig. S8A), while other cells Panc-1, PaTu, A-2058, and BLM were to some extent inhibited at concentrations of 1-5 μM (*SI Appendix*, Fig. S8B-E).

### Trafficking and compartmentalization of FP-5 in U-2OS cells

The ability of probe FP-5 to label cells effectively was monitored under various conditions and compared TF-Chol. When DMSO solutions of the probes were directly added to the cultivation medium containing 10% FCS, FP-5 labeled cells much faster than TF-Chol; intensive FP-5 fluorescence became apparent within 10-30 min while for TF-Chol, fluorescence was observed only at 24 h (Fig. 2A). Results from labeling of other cell lines with FP-5 (Raw 264.7, CHO-K1, MCF-7, and IEC-6) is available in *SI Appendix*, Fig. S9. When the cells were cultivated in lipoprotein-deficient serum (5% LPDS), labeling with both probes was enhanced and accelerated; FP-5 staining became apparent within 5-10 min and TF-Chol after 6 h of incubation (*SI Appendix*, Fig. S10A). Moreover, when cells were shortly pulsed by complexes of probes with the non-specific cholesterol carrier methyl β-cyclodextrin (MβCD) (molar ratio 1:10) the fluorescence of both probes was detectable on the cell surface immediately after the pulse (*SI Appendix*, Fig. S10B). FP-5, however, displayed more intensive fluorescence than TF-Chol in spite of the 10 times lower concentration used.

Time lapse microscopic images recorded during and after pulse confirmed very strong association of FP-5 with the plasma membrane (*SI Appendix*, 5. Movie Legend and Movie 1). Within 10 minutes the fluorescent signal appeared on intracellular membranes (*SI Appendix*, 5. Movie Legend, Movie 2), followed by increasing accumulation in the endo/lysosomal compartment within 0.5-2 h (*SI Appendix*, 5. Movie Legend, Movie 3). Co-localization studies with organelle specific probes confirmed association of FP-5 with endoplasmic reticulum (overlap with ER Tracker Red) after a 10 min chase (Fig. 2B) and increasing accumulation in the endo/lysosomal compartment (overlap with LysoTracker Red) after a 2-6 h chase (Fig. 2C). In some cases a slight transient signal appeared also in mitochondria (*SI Appendix*, Fig. S11A). A similar distribution, albeit with slower kinetics was observed with the TF-Chol probe (*SI Appendix*, Fig. S11B).

To find out the differences and similarities between the labeling of FP-5 and of dehydroergosterol (DHE), which is structurally close to cholesterol, we performed co-localization experiments. Importantly, pulse/chase experiments employing complexes MβCD/DHE together with MβCD/ FP-5 revealed a good co-localization between DHE and FP-5 shortly after pulse (*SI Appendix*, Fig. S11C, top right panel). However, after longer incubation (>30 min) FP-5 fluorescence concentrated gradually in lysosomes (see also Fig. 2C), whereas DHE was, even after 24 h incubation, localised diffusely in the cytoplasm (*SI Appendix*, Fig. S11C, low right panel). Thus, FP-5 in comparison to DHE showed a much faster redistribution into the lysosomal compartment.

### Heterocyclic sterol probes can detect cholesterol trafficking disorders

To model the situation in cholesterol trafficking disorders, we used an inhibitor of cholesterol transport U18666 and two model cell lines of human fibroblasts (GM03123E and GM18436) containing different mutations in the cholesterol transporter NPC1. Cells treated for 24-48 h with U18666A were labeled with FP-5, fixed and stained with filipin. Filipin fluorescence co-localized with FP-5 staining (Fig. 3A). FP-5 staining of NPC1 fibroblasts revealed intracellular punctuate structures in much higher abundance than in wild type human fibroblasts (HDFa) (Fig. 3B). These structures were also stained with filipin (Fig. 3C) and were identified as lysosomes according to co-labeling with Rhodamine dextran and LysoTracker Red (*SI Appendix*, Fig. S12). Intensive staining of lysosomal structures in NPC1 fibroblasts, with their accumulated cholesterol, was achieved faster with the FP-5 probe (within 2-6 h) than with TF-Chol (within 24 h) as shown in Fig. 3D. Likewise, other probes (FP-2, FP-6, FP-7, FP-8, and FP-10) exhibited ability for effective and fast labeling of cholesterol rich lysosomes in NPC1 cells (*SI Appendix*, Fig. S13A). These results show the potential of heterocyclic sterol probes for detecting cholesterol trafficking disorders.

**Fig. 1.**
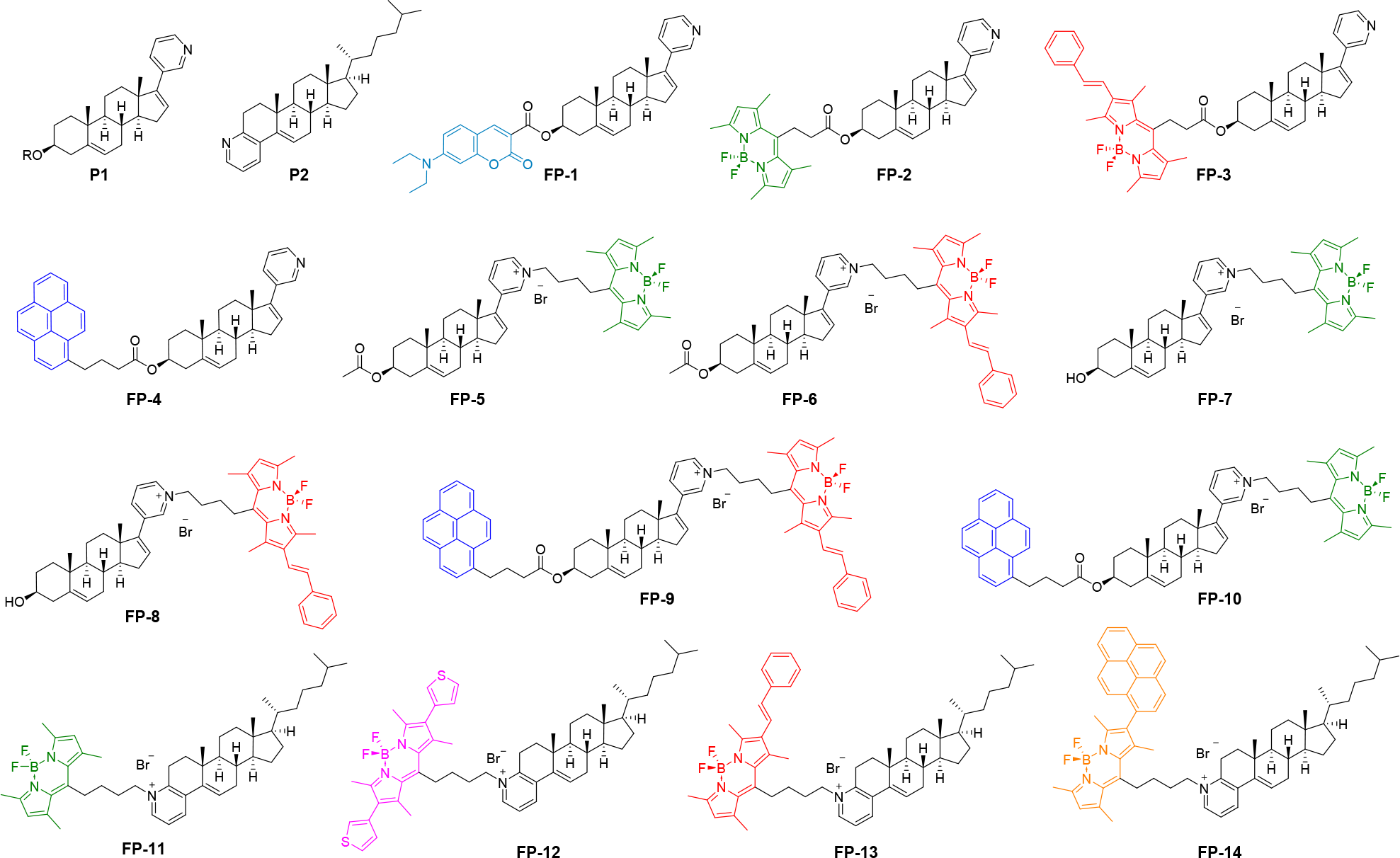
Heterocyclic precursors (P1 and P2) and synthetic probes (FP-1 - FP-14) for sterol sensing. R=Ac, H

## Discussion

Our study shows that heterocyclic sterol probes are suitable tools for cholesterol labeling, fulfilling the main requirements for usage, namely selective recognition, solubility, stability and good cellular uptake.

Generally, molecular recognition in biological systems relies on the existence of specific attractive interactions between two partner molecules. Structure-based drug or probe design seeks to identify and optimize such interactions between ligands and their target molecules. These interactions are given by their three-dimensional structures. The optimization process requires knowledge about the interaction geometries and approximate affinity contributions of attractive interactions, which can be gleaned from crystal structure and associated affinity data.

Since crystallographic data of probe-cholesterol complexes are not available in our hands, we believe that an acquired and unique cocrystal structure (*SI Appendix*, Figs. S4, S5) of the model sterol (in our case a cholesterol catabolite-lithocholic acid) with the heterocyclic structural model of our probe precursor (P1, abiraterone acetate) may indicate a possible mechanism of molecular recognition under studied conditions.

The probe structure was designed to provide a binding site for the steroid in question, in combination with pyridinium salt, ensuring sufficient aqueous solubility. Verification of our design validity came from spectroscopic studies as well as from X-ray crystallography. The van der Waals interactions were identified as main driving forces for crystal formation in aqueous environment, as cocrystal was obtained from methanol – water solution by slow evaporation. We assume that a similar binding mode takes place between at least some of our probes and sterol targets (see *SI Appendix*, Figs. S4 and S5). The rational design of probes used a combination of a) lipophilic part (steroid skeleton) with b) quaternary ammonium salt (giving expected selectivity and solubility) and c) fluorescent reporter, thereby creating unique fluorescent probes (*SI Appendix*, Figs. S1-S3). The quarternary pyridinium group was a point for fluorophore attachment. Importantly, the pyridyl group on the D ring (precursor P1, probes FP-1 – FP-8) seems to favour cellular uptake of probes in contrast to fused pyridine group on the A ring (precursor P2, probes FP-11–FP-14). The attachment of fluorescent groups on both sites of P1 (*via* 3-OH and pyridyl) resulted in insignificant cell fluorescence (FP-9) or aggregation of probe with slow cellular penetrance and fluorescence after 8-24 h incubation (FP-10) (Table 1, *SI Appendix*, Fig. S7). The variability in cellular uptake of probes, based on the same precursors, reflects the influence of other factors, such as the type of attached fluorophore and the presence of either acetyl or hydroxy groups on ring A. The probes FP-5 and FP-7 with BODIPY displayed fast and bright cellular fluorescence, while further extension of the fluorophore (increasing size) to generate a red BODIPY (FP-6, FP-8) led to slower uptake and lower fluorescence. The presence of acetyl or hydroxyl group in FP-5 and FP-7, respectively, affects lipophilicity and consequently the kinetics and trafficking of probes. Generally, we observed a correlation between lipophilicity of compounds and their cellular uptake, e.g. highly lipophilic FP-3, FP-4, FP-9, and FP-14 probes did not enter cells, while less lipophilic FP-5 – FP-8 effectively labeled cells (see *SI Appendix*, Table 1, Table S1, Fig. S7). Notably, FP-5 displayed a durable association with cellular membrane structures whereas FP-7, with its hydroxyl group, displayed a markedly shorter association with plasmatic and intracellular membranes, and profound labeling of lysosomes (*SI Appendix*, Fig. S7). Thus, it can be concluded that a delicate tuning of lipophilicity, which can be increased by acetyl group is required for optimal probe properties. On the other hand, fast labeling of lysosomes by FP-7 may be an advantage for detecting lysosomal storage disorders, as we observed very intensive staining of NPC1 fibroblasts using this probe within 2 h (*SI Appendix*, Fig. S13B).

The favourable properties of FP-5 for monitoring of sterol trafficking, emerged from comparison with the commercial probes, TF-Chol and DHE. The application of FP-5 directly to the growth medium, resulted in the effective labeling of cells within 30 min, while for TF-Chol, intracellular fluorescence was only visible after 24 h incubation (Fig. 2A). This demonstrates that FP-5 diffuses from solvent to cells quickly, whereas the hydrophobic TF-Chol, having a propensity to aggregate in an aqueous environment, equilibrates more slowly with intracellular membranes (27, 28). Similar aggregates formation and slow labeling occurred also with some of our probes (FP-2, 10, 11, and 12). Moreover, it is known that TF-Chol is readily effluxed from cells and prolonged incubation in the presence of efflux acceptors like apolipoprotein A-I may significantly reduce the signal intensity (27). Accordingly, in their absence in medium supplemented with the lipoprotein-deprived serum (LPDS), we observed a significant fluorescence already after 6 h (*SI Appendix*, Fig. S10A). A major improvement of TF-Chol signal was achieved using the artificial carrier MβCD in LPDS conditions. This labeling method yielded a robust and uniform plasma membrane signal, which is consistent with reports of others (19, 27, 29). These conditions also increased and accelerated FP-5 signal (*SI Appendix,* Fig. S10B). Similarly, a DHE signal was achieved only when complexed with MβCD (29). It can be summarized that FP-5 shows better uptake than commercial probes, which mostly require artificial carriers and the absence of efflux acceptors in order to effectively label cells. In spite of the fact that an extensive comparative study with other fluorescently labeled cholesterol analogs and their performance in various cellular assays, similar to work of Sezgin (18), is still needed, the applicability of FP-5 is evident.

Time lapse experiments revealed a robust and dynamic plasma membrane signal of FP-5, during and after the labeling pulse, with fast movement along membranes and extensive influx and efflux, (*SI Appendix*, Movie 1-3). After the pulse, an increasing intracellular signal displayed transient co-localization with ER Tracker. ER is the organelle where most steps of cholesterol synthesis take place, but cholesterol is rapidly transferred from the ER to other organelles *via* vesicular trafficking and nonvesicular pathways (4). There are reports showing existence of endoplasmic reticulum (ER)–plasma membrane (PM) junctions as contact sites between the ER and the PM (30, 31). These membrane contact sites (MCSs) are domains where two membranes come to close proximity, typically less than 30 nm, that favor exchange between the two organelles. They are established and maintained in durable or transient states by tethering structures, which keep the two membranes in proximity without fusion. The ER extensive network is involved in the most MCSs within the cell, including mitochondria, lysosomes, lipid droplets, Golgi apparatus, and endosomes (32, 33). It is possible that the FP-5 transport from PM into ER and subsequently to lysosomes is mediated through MSC by the nonvesicular pathway. Such an interpretation is supported by reports describing MCSs involvement in PM-ER sterol transport (34, 35). However, the involvement of vesicular transport cannot be ruled out.

Fast labeling of NPC1 fibroblasts with defective cholesterol trafficking (36–38) in normal cultivation medium is another advantage of FP-5 (Fig. 4B). A strong and uniform signal in the lysosomal compartment appeared within 2 h, while a TF-Chol signal took 24 h to develop (Fig. 3D). Notably, when NPC1 cells were labeled with FP-7, an early bright signal appeared in mitochondria (30 min) before prevailing in the endolysosomal compartment (cca 2 h) (*SI Appendix*, Figs. S13B). A recent report indicates the possibility of lysosomal-mitochondrial liaisons leading to accumulation of specific lipids and cholesterol in mitochondria. Such accumulation results in mitochondrial dysfunction and defective antioxidant defence, contributing to Niemann-Pick disease progression (39). In line with this, our observation suggests an attractive possibility that FP-7 might be a promising tool for studying lysosomal and mitochondrial interactions and sterol trafficking in this disease. In addition, some other heterocyclic sterol probes (FP-2, FP-6, FP-8 and FP-10) label NPC1 cells effectively within 6 h and hence can be used for detection of lysosomal storage disorders (*SI Appendix,* Fig. S13A).

In summary, this study demonstrates FP-5 as a unique probe providing uniform and fast labeling of plasmatic and intracellular membranes, transferring later to their ultimate destination, the lysosomes. Fig. 4 summarizes the schematic visualization of dynamic sterol transport and intracellular trafficking in a U-2 OS cell (A), and pathological accumulation in a NPC1 fibroblast (B). The analogical scheme for TF-Chol signal distribution with much slower kinetics than for FP-5 is shown in *SI Appendix*, Fig. S14. A preliminary comparison with two commercial probes, TF-Chol and DHE highlights advantages of the described novel class of heterocyclic sterol probes, applicable for monitoring of sterol trafficking and its pathologies.

**Fig. 2.**
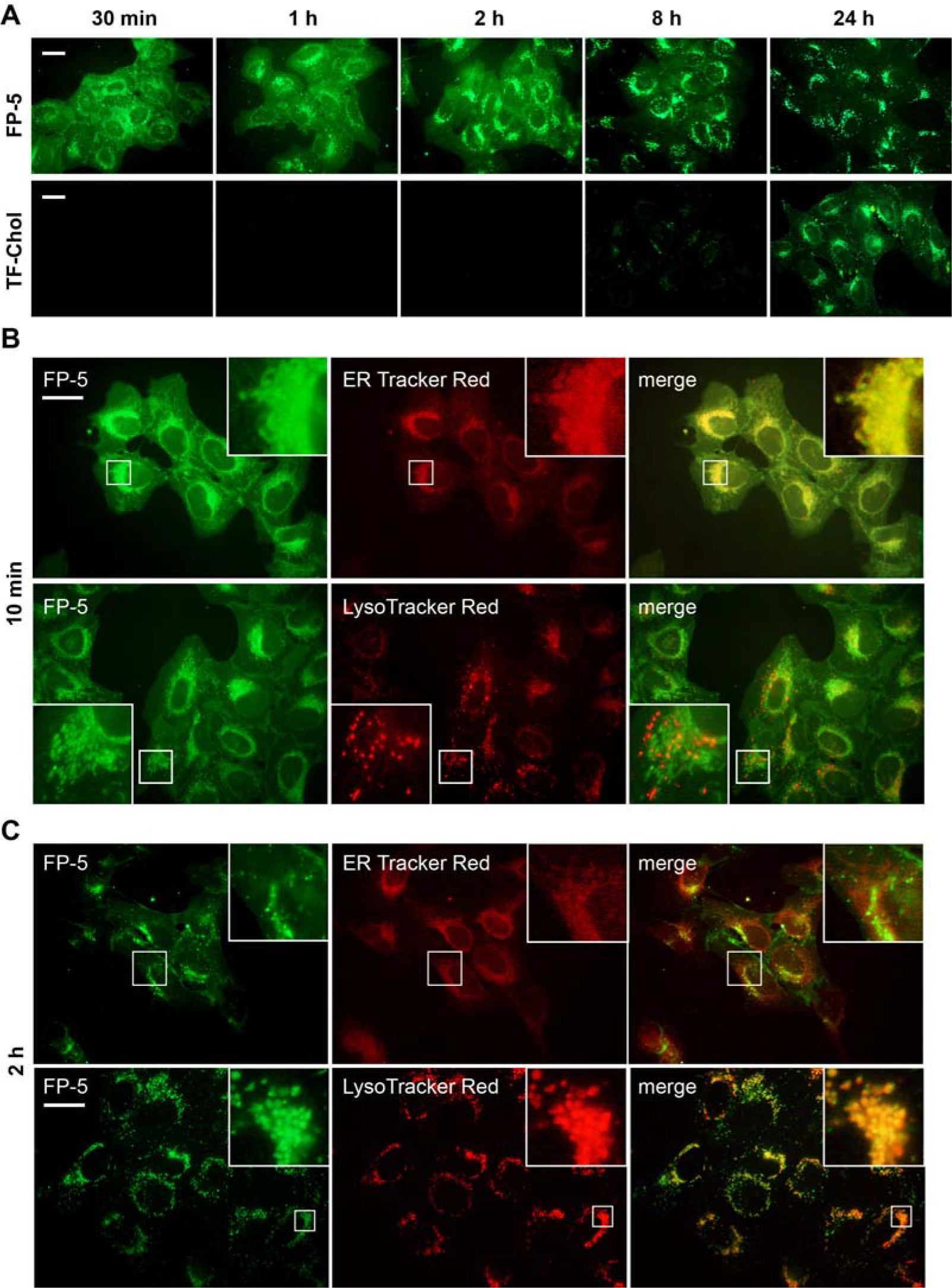
The kinetics and localization of intracellular fluorescence of FP-5 and TF-Chol in U-2 OS cells. (**A**) Probes in DMSO solution were directly added to cultivation medium with 10% FCS at final concentration 0.5 μM and live fluorescence was recorded at indicated time points. Scale bar represents 10 μM. (**B-C**) Co-localization of FP-5 and organelle specific probes. Cells exposed to pulse with complex FP-5/MβCD (1 μg/ml) were chased for 10 min or 2 h, co-labeled with ER Tracker Red or LysoTracker Red and examined. Expansions of the regions indicated by the white boxes are shown on the upper right side or low left side. Localization of TF-Chol is included in *SI Appendix*, Fig. S15).

**Fig. 3.**
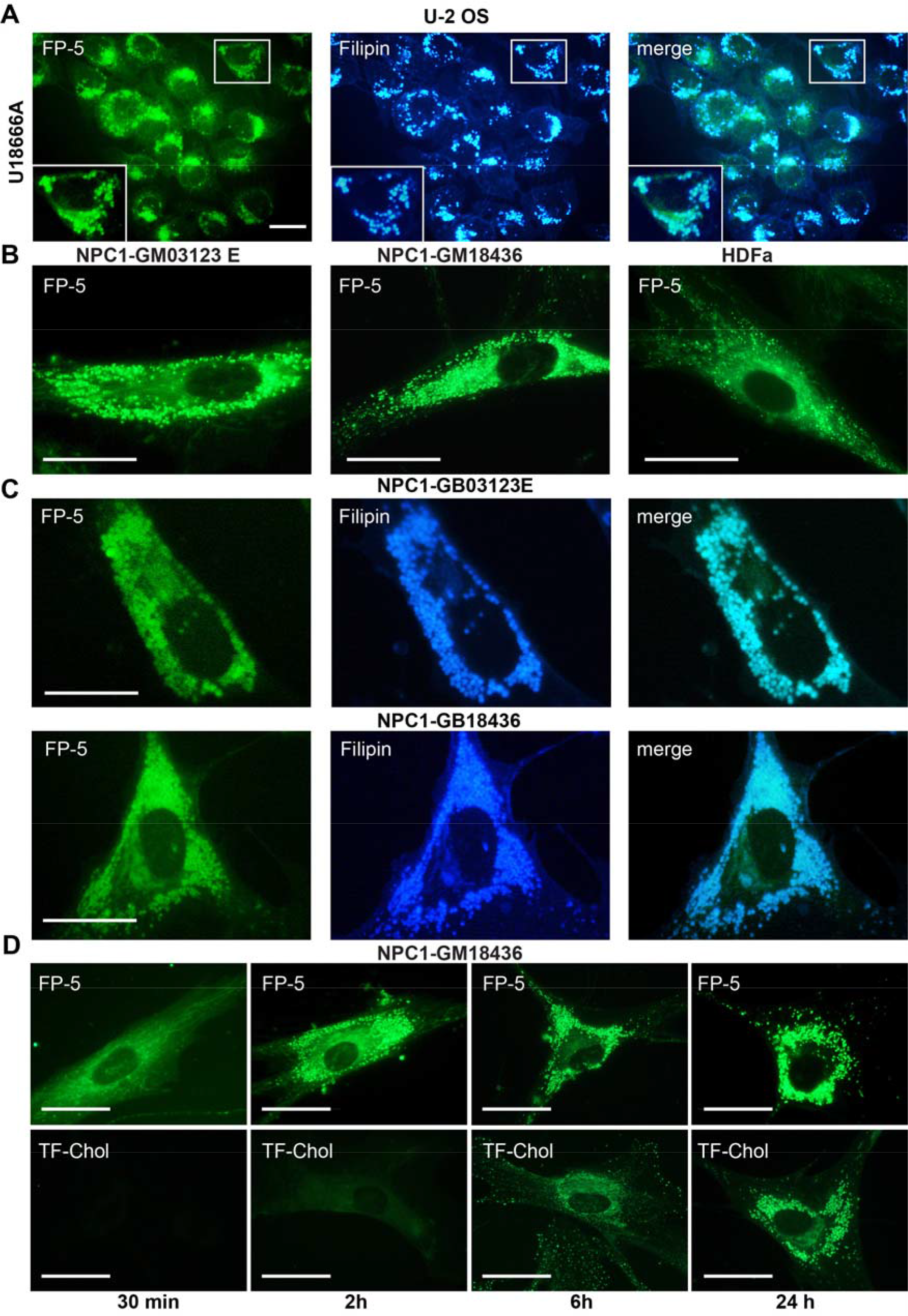
FP-5 fluorescence in cells with abnormal content of cholesterol. (A) Cholesterol transport in U-2 OS was inhibited by inhibitor U18666A (1 μg/ml) for 48 h and then cells were labeled with FP-5 (200 nM) for additional 24 h, fixed and stained with filipin (50 μg/ml). Expansion of the region indicated by the white box is shown on the low left side. (B) Human fibroblasts carrying mutations in NPC1 cholesterol transporter (clones GB03123E, GB18436) and control normal human fibroblasts (HDFa) were labeled with FP-5 (200 nM) for 6 h and examined. (C) Co-localization of FP-5 and filipin staining in mutant cell clones. (D) Differential kinetics of FP-5 and TF-Chol lysosomal labeling in NPC-GM18436 fibroblasts. Cells were incubated with FP-5 (200 nM) and TF-Chol (1μM) for indicated times in medium containing 5% LPDS and imaged live. Scale bar represents 10 μM.

**Fig. 4.**
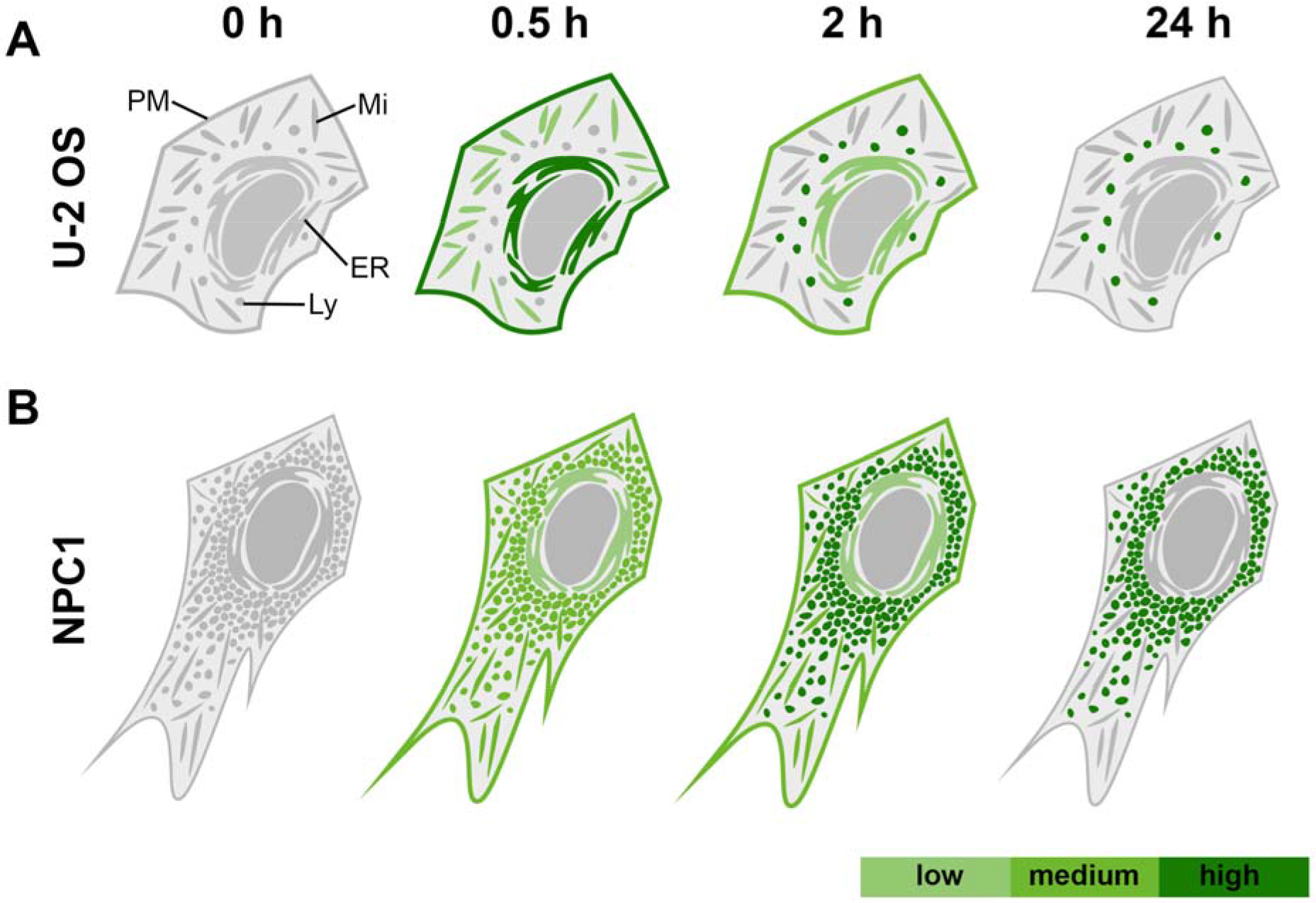
Schematic distribution of FP-5 signal in (A) U2-OS cells and (B) in Niemann-Pick fibroblasts following direct addition of FP-5 solution to cultivation medium. The FP-5 signal in U-2 OS cells progresses from plasmatic membrane to ER (time 0.5 h) and accumulates in lysosomes (2-24 h). In NPC1 fibroblasts with accumulated cholesterol, FP-5 signal appears in lysosomes quickly and intensifies within 0.5-2 h.

**Table 1.**
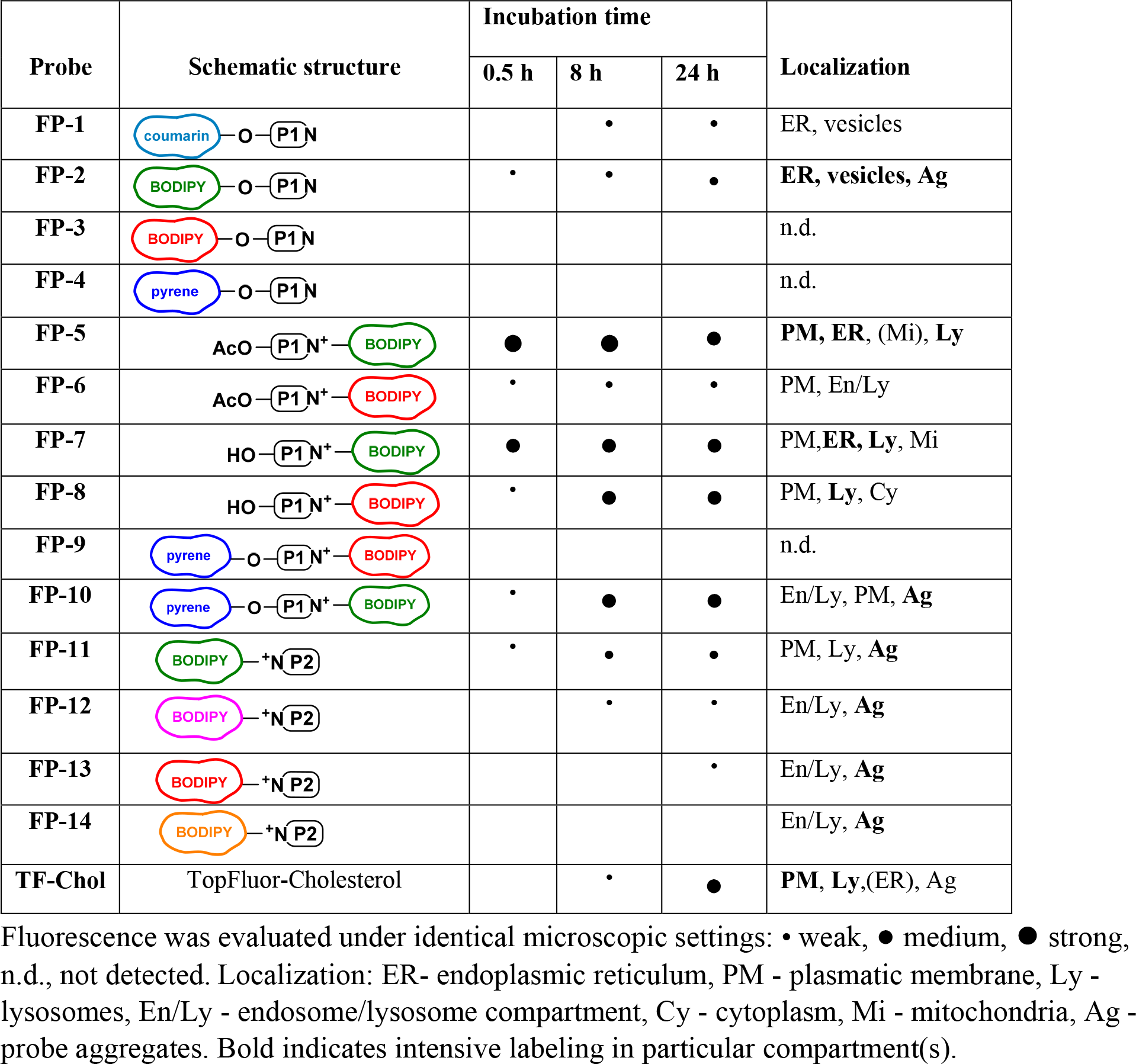
Intensity of intracellular fluorescence of novel sterol sensing probes.

## Materials and Methods

### Reagents and Materials

Solvents were purchased from PENTA, and steroids from Steraloids. The purchased material was used without further purification or distillation. TopFluor-Cholesterol (TF-Chol) was from Avanti Polar Lipids, Inc., Rhodamine-Dextran, LysoTracker^®^ Red DND-99, and ER-Tracker™ Red were from Molecular Probes (Life Technologies). Dehydroergosterol (DHE), methyl-β-cyclodextrin, LPDS (lipoprotein deficient serum) and Filipin were from Sigma, inhibitor U-18666A from ENZO Life Sciences and media RMPI, EMEM and supplements were from Life Technologies.

### Chemical Synthesis and compound characterization

Synthesis of compounds and their characterization is included in *SI Appendix*, 1. Chemistry.

### Cell Lines and Cell Culture

U-2 OS cells (obtained from ATCC) were cultivated in RPMI 1640 medium supplemented with 10% FBS (Life Technologies), sodium pyruvate, 2 mM glutamine, penicillin and streptomycin (Sigma), 20 mM HEPES, and glucose (4mg/ml). NPC1 fibroblasts (obtained from Coriell Repository) were maintained in EMEM medium supplemented with NEAA (nonessential amino acids) (Life Technologies), 2 mM glutamine, penicillin and streptomycin (Sigma), and 15% FCS. HDFa fibroblasts were grown in Medium 106 with Low Serum Growth Supplement (LSGS) as recommended by manufacturer (Life Technologies).

### Labeling of cells with fluorescent probes and organelle markers

Solutions of probes were prepared in DMSO and applied to cultivation medium (200 nM – 1 μM final concentration) supplemented with either 10 % FBS or 5% LPDS. To facilitate cellular uptake, probes were complexed with methyl-β-cyclodextrin (MβCD) at a molar ratio of 1:10 (probe:cyclodextrin) as described before for BODIPY-cholesterol (27), sonicated 2 × 3 min and centrifuged for 5 min. The complex was applied to cells for 2 min at room temperature at a concentration 0.5-2 μg/mL of FP-5 and 20 Fg/mL of TF-Chol.

For double-labeling cells with DHE and FP-5 cholesterol analogs, analogs were loaded on methyl-β-cyclodextrin (29). The concentrations of DHE and FP-5 used for preparation of the cyclodextrin (CD) complexes were 3 mM and 1.5 μM, respectively. Thus, DHE was in 2000-fold excess to FP-5. Cells were labeled with DHE/FP-5/MβCD for 2 min at room temperature, washed and chased for 10 min, 4 and 24 h at 37 °C.

For co-localization studies, cells were incubated with probes and subsequently loaded with organelle markers LysoTracker™ Red DND-99 (80 nM), or ER-Tracker™ Red (1 μM) (Molecular Probes) for 30 min at 37 °C in complete medium. Rhodamine-dextran (1 mg/mL) was supplied overnight in normal growth medium. For Filipin co-staining, cells were, after labeling with probes, fixed with 3% paraformaldehyde and then labeled with filipin (50 μg/ml) for 30 min (15).

### Fluorescence Microscopy

Cells grown on coverslips in 35-mm Petri dishes were incubated with the corresponding probe for indicated time in FluoroBright™ DMEM medium without phenol red, subsequently washed and observed alive using a fluorescence microscope DM IRB (Leica) with filter cube I3 (excitation filter BP 450–490 nm and long pass filter LP 515 nm for emission) for green fluorescence, filter cube N2.1 (excitation filter BP 515-560 nm and long pass filter LP 590 nm for emission) for red fluorescence, and filter cube A (excitation filter BP 340–380 nm and long pass filter LP 425 nm for emission) for blue fluorescence. The fluorescence images were acquired by a DFC 480 camera using a 63× oil immersion objective.

### Time-lapse microscopy

For the time lapse experiments, U-2 OS cells were plated on 35 mm glass bottom dishes (In Vitro Scientific) coated with poly-L-lysine. The dish was placed within a humidified chamber (37 °C, 5% CO2) on the microscope stage for 30 min. Cells were then pulsed with FP-5/M/CD for 2 min (final concentration of FP-5 was 0.5 μM), washed and then the fluorescence signal was acquired. The acquisition was performed on an OMX Delta Vision microscope in wide-field mode; microscope settings: objective 60×/1.42NA PlanApo N, excitation filter 477/32, emission filter 528/48. Images were acquired on a PCO.EDGE sCMOS camera, readout speeds 95 MHz. Frame rate interval for acquisition was 10 seconds.

## ACKNOWLEDGMENTS

The authors would like to thank Trevor Epp for review of the manuscript and Tereza Mikulášová for drawing schematic figures. This work was supported by The Czech Science Foundation (grant 17-02836S), by the programs of Ministry of Education Youth and Sports of the Czech Republic (MEYS) LO1220, LM2015063, and Operational Program Prague Competitiveness financed from the European Regional Development Fund (ERDF OPPC) (CZ.2.16/3.1.00/24510). We acknowledge the Light Microscopy Core Facility, IMG CAS, Prague, Czech Republic supported by MEYS (LM2015062), OPPC (CZ.2.16/3.1.00/21547) and (LO1419) for their support presented herein.

## Author contributions

J.K. performed research and wrote the paper, M.J. contributed new reagents or analytic tools, L.K., B.D., Z.R., M.D., P.D., P.B. analyzed data, I.N., M.K. performed research, P.B. and V.K. designed research.

